# Genomic characterization of the virulence-associated pyomelanin biosynthetic pathway in pigment-producer strains from the pandemic *Acinetobacter baumannii* IC-5

**DOI:** 10.1101/2020.07.10.197806

**Authors:** Érica Fonseca, Fernanda Freitas, Raquel Caldart, Sérgio Morgado, Ana Carolina Vicente

## Abstract

**BACKGROUND:** *Acinetobacter baumannii* outbreaks have been associated with pandemic International Clones (ICs), however the virulence factors involved with their pathogenicity are sparsely understood.

**OBJECTIVES:** This study aimed to characterize the reddish-brown pigment produced by *A. baumannii* strains, and to determine its biosynthetic pathway by genomic approaches.

**METHODS:** Pigment characterization was conducted by phenotypic tests in different growth conditions. The clonal relationship among *A. baumannii* was obtained by PFGE and MLST and antimicrobial susceptibility test was performed by Disc-Diffusion method. The genome of one representative strain was obtained for characterization of genes involved with pigment production.

**FINDINGS:** The virulence-associated pyomelanin was the pigment produced by *A. baumannii*. These strains were extensively drug resistant and belonged to the IC-5/ST79. Genomic approaches revealed that the pyomelanin biosynthetic pathway in *A. baumannii* presented a particular architecture concerning the peripheral (*tyrB, phhB* and *hpd*) and central (*hmgB, hmgC* and *hmgR*) metabolic pathway genes. The identification of a distant HmgA homologue, probably without dioxygenase activity, could explain pyomelanin production. Virulence determinants involved with adherence (*csuA/BABCDE* and a T5bSS-carrying genomic island), and iron uptake (*basABCDEFGHIJ, bauABCDEF* and *barAB*) were also characterized.

**MAIN CLONCLUSION:** The pyomelanin production together with other virulence determinants could play a role in *A. baumannii* pathogenicity.

---

*Acinetobacter baumannii* is one of the most relevant pathogens associated with nosocomial infections that presents the long-term ability to survive on inanimate surfaces, contributing to national and international clonal dissemination.^(1)^ The *A. baumannii* outbreaks have been associated with high-risk pandemic lineages, named International Clones (ICs), characterized by a high capacity to persist in clinical environments and by presenting a broad antimicrobial resistance profile.^(2,3)^ However, in spite of *A. baumannii* association with nosocomial and persistent infections, the role of virulence factors in its pathogenesis remains largely obscure. This virulence has been associated with features that enhance its persistence, such as increased adherence, resistance to dissection, biofilm formation, production of capsule and iron uptake.^(4–6)^

In bacteria, the production of pigments, as melanins, have been linked with virulence and pathogenicity. Melanins are a black-brown and yellow-red pigments derived from the oxidation of different phenolic compounds.^(7)^ Depending on the biosynthesis pathway, melanin may be given a different designation, such as pyomelanin, which is a reddish-brown pigment resulted from tyrosine (Tyr) or phenylalanine (Phe) through the accumulation of homogentisic acid (HGA).^(8)^ This pigment provides protection against oxidative stress and contribute to invasiveness and persistence by enhancing bacterial surface attachment and biofilm formation, extracellular electron transfer, resistance to heavy-metals and iron reduction/acquisition, induction of virulence factor expression, contributing to the adaptive response to environmental stress.^(9,10)^ The pyomelanin production results from a defect in the catabolism pathway. *Pseudomonas putida* metabolizes Phe and Tyr through a peripheral pathway, regulated by the σ^54^-dependent transcriptional activator PhhR, involving hydroxylation of Phe to Tyr by PhhAB, conversion of Tyr into 4-hydroxyphenylpyruvate by TyrB, and formation of HGA by Hpd as the central intermediate. HGA is then catabolized by a central catabolic pathway that involves the homogentisate dioxygenase (HmgA), fumarylacetoacetate hydrolase (HmgB), and maleylacetoacetate isomerase (HmgC), finally yielding fumarate and acetoacetate.^(11)^ Mutations or deletions that result in loss of HmgA function, as well as overexpression of *hmgR*, a *hmgA* repressor from the Tet^R^ family, lead to an accumulation of HGA.^(12,13)^ The accumulated HGA is then secreted from the cell via the HatABCDE ABC transporter, where it auto-oxidizes, and self-polymerizes to form pyomelanin.^(14)^ The production of this pigment is quite common in species such as *Legionella, Vibrio cholera* and *Pseudomonas sp*.^(9–13)^ However, pyomelanin production in *A. baumannii* was only reported in a lineage causing nosocomial infections in Rio de Janeiro city, Brazil, in 2010.^(15)^

This study reports the occurrence of persistent *A. baumannii* strains producing a brown diffusible pigment resembling the pyomelanin, which caused an outbreak in a hospital of the Amazon Basin, Brazil. Based on whole genome analyses of a representative strain, we characterized the biosynthetic pathway involved with the production of this pigment, which could contribute to the virulence of this *A. baumannii* lineage.

## MATERIAL AND METHODS

### Clinical data, bacterial strains and antimicrobial susceptibility test

From October, 2016 to April, 2018, 12 *A. baumannii* producing a brown diffusible pigment were recovered from inpatients hospitalized in different wards of the General Hospital of Roraima (HGR), placed in Boa Vista, a city embedded in the Amazon Basin, Brazil (TABLE). Species identification was determined by the automated VITEK2, and confirmed to be *A. baumannii* by sequencing the 16S rRNA and the *bla*_OXA-51_ genes.

**TABLE.** Clinical and genotypic features of the XDR pyomelanogenic *A. baumannii* strains.

|  | **Isolation date** | **PFGE** | **MLST (IC)** | **Ward** | **Clinical specimen** |
| --- | --- | --- | --- | --- | --- |
| AB4353 | Oct/21/16 | A | ST79 (IC-5) | ICU | Catheter tip |
| AB77 | Jan/05/17 | A | ST79 (IC-5) | others hospital wards | Wound secretion |
| AB2299 | Feb/12/17 | A | ST79 (IC-5) | Emergency | Bronchial aspirate |
| AB1077 | Mar/08/17 | A | ST79 (IC-5) | ICU | Tracheal secretion |
| AB1113 | Mar/08/17 | A | ST79 (IC-5) | ICU | CRL |
| AB81 | Oct/02/17 | A1 | ST79 (IC-5) | ICU | Tracheal secretion |
| AB04-RR5 | Jan/01/18 | A1 | ST79 (IC-5) | ICU | Blood |
| AB28-RR5 | Jan/17/18 | A | ST79 (IC-5) | ICU | Tracheal secretion |
| AB41-RR4 | Jan/19/18 | A | ST79 (IC-5) | ICU | Tracheal secretion |
| AB51-RR5 | Jan/26/18 | A | ST79 (IC-5) | ICU | Tracheal secretion |
| AB04-RR6 | Apr/25/18 | A | ST79 (IC-5) | ICU | Catheter tip |
| AB05-RR6 | Apr/29/18 | A | ST79 (IC-5) | others hospital wards | Catheter tip |

CRL, Cephalorachidian liquid

The antimicrobial susceptibility test was determined by the disc-diffusion method, according to CLSI guidelines,^(16)^ for the following antibiotics: gentamicin, amikacin, tobramycin, imipenem, meropenem, doripenem, ciprofloxacin, ampicillin/sulbactam, piperacillin/tazobactam, ticarcillin/clavulanic acid, cefotaxime, ceftazidime, cefepime, trimethoprim/sulphamethoxazole, tetracycline and minocycline. The MIC of polymyxin B was assessed by the broth microdilution with antibiotic concentrations ranged from 0.1 μg/ml to 64 μg/ml. The current definition criteria for classifying *A. baumannii* antimicrobial resistance was applied.^(17)^

### Phenotypic characterization of brownish pigment produced by A. baumannii strains

The strains were grown overnight on Mueller-Hinton (MH) and trypticase soy agar (TSA) media plates at different temperatures (28°C, 35°C and 40°C) to verify the influence on pigment production. To investigate whether the pigment is the pyomelanin resulted from the tyrosine metabolism, the 12 pigment-producing *A. baumannii* strains were grown in a minimal medium (T-Medium),^(18)^ with the tyrosine and glutamate as the sole carbon sources. The pyomelanin-producing *A. baumannii* 456MDp,^(15)^ kindly provided by Dr. Beatriz M. Moreira, and the *A. baumannii* ATCC 19606 were also included in this test as positive and negative controls, respectively.

An additional test was performed to determine which tyrosine metabolic pathway was involved with the brown pigment production. Therefore, the effect of sulcotrione [2-(2-chloro-4-methane sulfonylbenzoyl)-1,3-cyclohexanedione)], an inhibitor of tyrosine metabolism via homogentisic acid,^(19)^ was evaluated by growing the isolates in the T-medium in the presence of different concentrations (2.5, 10, 15 and 20 mM) of sulcotrione.

### Determination of genetic relatedness of A. baumannii strains

The genetic relationship among the 12 pigment-producing *A. baumannii* strains and between these strains and the pyomelanin-producing *A. baumannii* 456MDp, previously identified in a hospital from Rio de Janeiro,^(15)^ were assessed by PFGE and MLST using the Pasteur and Oxford schemes (https://pubmlst.org/abaumannii/) available in the *A. baumannii* MLST website.

### Whole genome sequencing and genome annotation

The genome sequence of one representative pigment-producing strain (AB4353) were obtained with the Illumina HiSeq 2500 sequencer using Nextera XT paired-end run with a ~500-bp insert library at the High-Throughput Sequencing Platform of the Oswaldo Cruz Foundation (Fiocruz, Rio de Janeiro, Brazil). The quality of the reads was assessed with FASTQC and *de novo* assembling was performed with the SPAdes 3.5 assembler with default settings. Gene prediction and annotation were performed with RAST tool and Prokka software (https://github.com/tseemann/prokka). The resistome was assessed with the Comprehensive Antibiotic Resistance Database (CARD) (https://card.mcmaster.ca/). The mobilome and virulome were assessed with IslandViewer4 (https://www.pathogenomics.sfu.ca/islandviewer/) and VRprofile 2.0 (https://bioinfo-mmi.sjtu.edu.cn/VRprofile/) web servers, respectively. AB4353 genome sequence has been submitted to GenBank under accession no. JAAXKU000000000.1.

## RESULTS AND DISCUSSION

### Characterization of brownish-producer A. baumannii strains

The 12 clinical *A. baumannii* strains producing a brown diffusible pigment were phenotypically and genotypically characterized. All of them presented the extensively-drug resistant (XDR) phenotype, since they were susceptible only to polimixin B and tetracyclines.

All strains were able to produce the pigment on MH medium at all tested temperatures, with a more prominent production at higher temperatures (35°C and 40°C) (Fig. 1A, data shown for AB4353, AB1077, AB1113, AB41-RR4), as previously demonstrated.^(15,20)^ It was verified that production of pigment at higher temperatures was due to the induction of *melA* (*hpd*), which is responsible for the HGA synthesis,^(20)^ suggesting the role of this mechanism in the adaptive response to environmental stress. On the other hand, no pigment was observed on TSA medium.

**Fig. 1:**
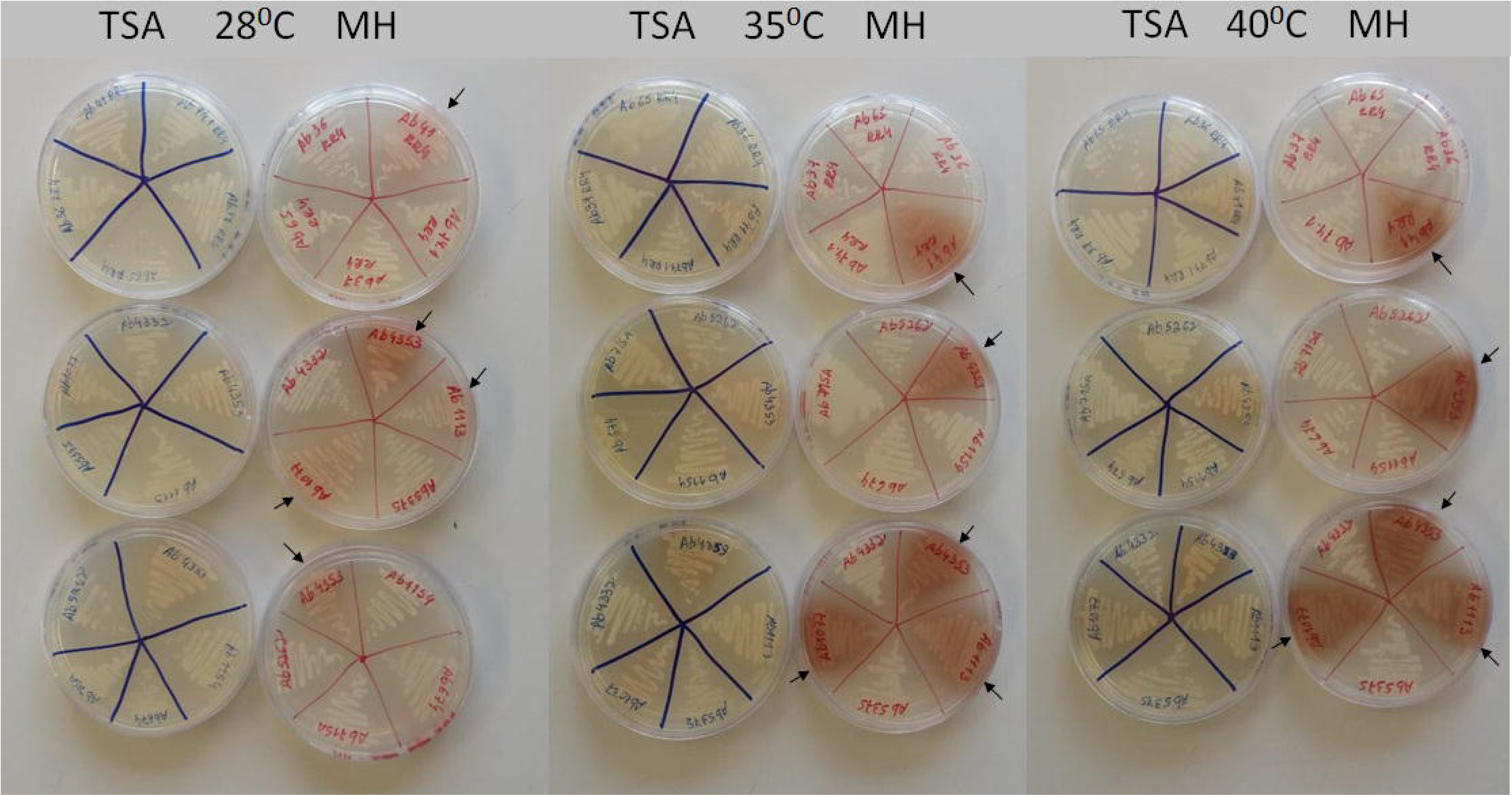
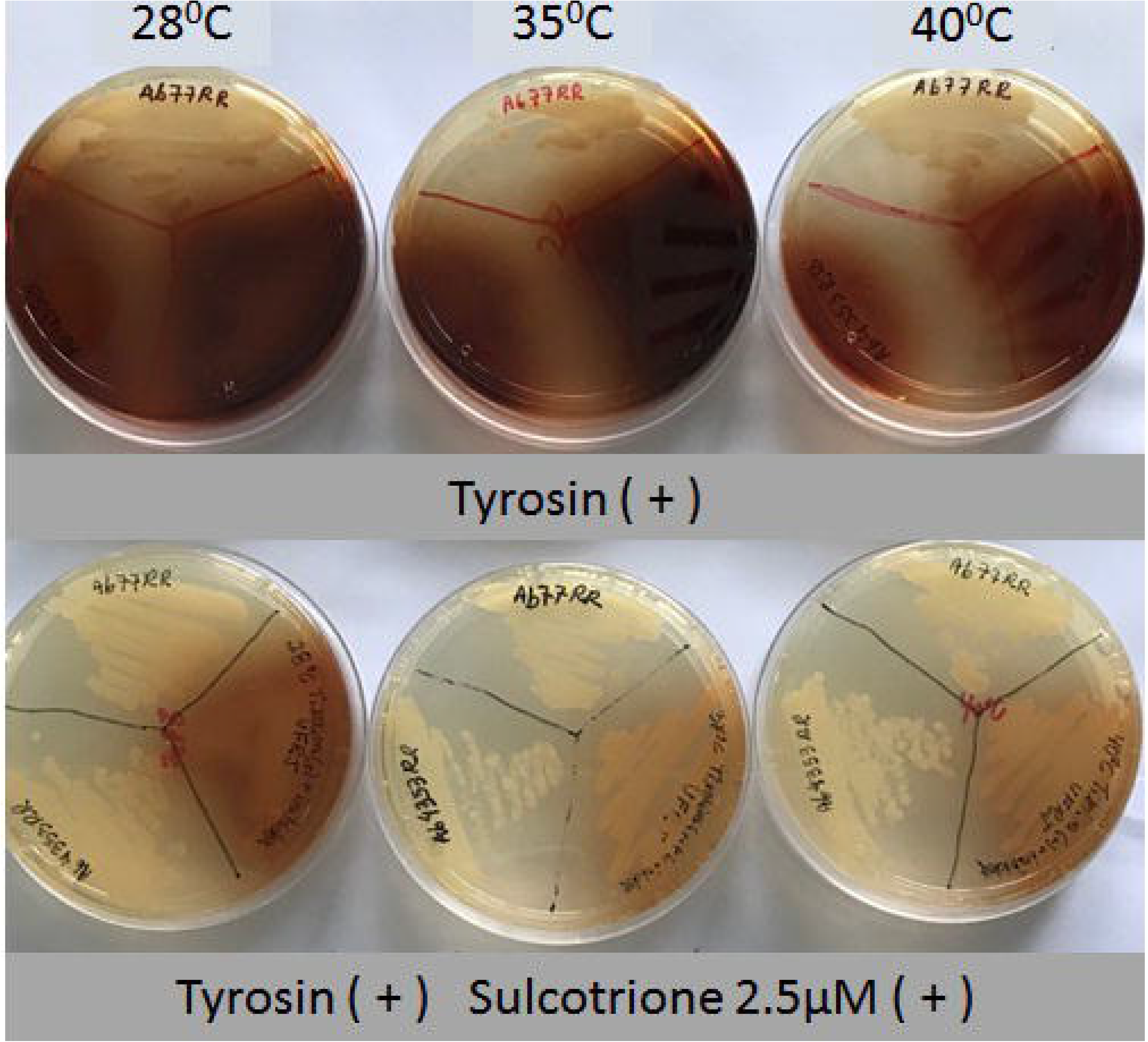
Pyomelanin production by the XDR *A. baumannii* strains. **(A)** The conditions and temperatures used in the test are shown. Arrows indicate the pyomelanogenic *A. baumannii* from this study chosen as representative strains for these tests (AB4353, AB1077, AB1113, AB41-RR4). Other *A. baumannii* strains were used as negative controls; **(B)** Production of pyomelanin at minimal T-medium in the presence and in the absence of sulcotrione inhibitor at different temperatures. The image shows the pigment production by two pyomelanogenic *A. baumannii* representative strains of this study (AB4353 and AB77) and by the 456MDp strain used as positive control.

Phenotypic tests to verify whether the brown pigment resulted from the tyrosine catabolic pathway revealed that, after 20h of incubation on T-medium in different temperatures, a brown diffusible pigment was observed in the 12 *A. baumanniii* and in the positive control 456MDp strain (Fig. 1B, data shown for AB4353, AB77 and 456MDp), while no pigment production was observed on the negative control ATCC 19606 (data not shown). Moreover, a significant reduction in the brown pigment production occurred on T-medium plus 2.5 μM of sulcotrione (Fig. 1B). Considering that this substance is an inhibitor of tyrosine metabolism via HGA pathway, and that pyomelanin production results from its accumulation and efflux, it could be inferred that this brownish pigment corresponded to pyomelanin.

### Genetic relatedness and epidemiology of pyomelanin-producer A. baumannii

The PFGE revealed that the 12 XDR pyomelanin-producing *A. baumannii* strains were clonally related (TABLE), however, no genetic relationship was observed between these strains and the pyomelanogenic *A. baumannii* 456 MDp strain recovered from a hospital in Rio de Janeiro in 2010.^(15)^ All 12 strains belonged to ST79^PAS^/ST758^OXF^, which corresponds to the high-risk pandemic International Clone V (IC-5), while the 456MDp strain belonged to ST1079^PAS^/ST1483^OXF^ already identified in China in 2015 (MLST metadata). The IC-5 is prevalent in clinical settings spread in Brazil and South America,^(21)^ however, the pigment production has never been highlighted as a phenotypic trait of this IC neither in Brazil nor in other continents. Interestingly, this lineage has persisted in HGR for more than one year (19 months), which could be resulted from an increased adaptive fitness.

### The resistome of AB4353

Resistome prediction analyses of AB4353 revealed the presence of several genes, conferring resistance to aminoglycosides (*aac(6’)-Ian, aac(3’)-IIe, aph(3”)-Ib, aph(6’)-Id*), chloramphenicol (*cmlA*), sulfonamide (*su11*), β-lactams (*bla*_TEM-1b_ and *bla*_OXA-65_) and carbapenems (*bla*_OXA-23_), corroborating the observed XDR phenotype.

### Virulome of pyomelanin-producer AB4353 strain: adherence, iron uptake and desiccation tolerance

Bacterial adherence constitutes an essential step in the colonization process. *In silico* analysis of AB4353 genome revealed the presence of a 18 kb adherence-related genomic island previously identified in AbH120-A2, an *A. baumannii* strain with a remarkable adherence ability responsible for a large nosocomial outbreak in Spain from 2006 to 2008.^(22)^ This island carried the Type Vb secretion system from the two-partner System (TPS) family composed by TpsA (AbFhaB) and TpsB (AbFhaC), a large exoprotein involved with Heme utilization and adhesion and its translocator channel, respectively. This adhesion-related secretion system had already been found in other Gram-negative bacteria, and it is considered one of the main virulence factors in *Bordetella pertussis*.^(23)^ Interestingly, the AbH120-A2 (2006-2008) from Spain and AB4353 (2017) from Brazil belongs to the IC-5 (ST79), demonstrating the increased adaptive fitness and the remarkable spread potential of this lineage. Such adaptation could be due to the presence of this adhesion-related island, among other factors, which has been probably contributing to IC-5 persistence in clinical settings worldwide for, at least, 10 years. Additionally, other determinants associated with biofilm formation and adherence phenotypes were also identified in AB4353, such as the biofilm-associated protein (Bap) and the CsuA/BABCDE usher-chaperone system.^(24–26)^

The iron uptake capacity has been considered an important component for bacterial growth and survival under iron-limiting conditions found in host environment, also contributing to pathogenicity. The AB4353 harbour the siderophore Acinetobactin operon identical to that found in the ATCC 19606^T^, composed by *basABCDEFGHIJ, bauABCDEF and barAB* genes, involved with biosynthesis, utilization and siderophore release, respectively.^(27)^

Desiccation tolerance contributes to the remarkable persistence character of *A. baumannii*, allowing it to become a successful pathogen in the nosocomial environment. The two-component System BfmRS is directly involved with the production of the desiccation resistance phenotype in this species.^(28)^ Two residues in BfmR, Leu230 and Thr85, are crucial to the BfmR activity and the control of stress responses, which protect *A. baumannii* cells during desiccation. The deduced BfmR from AB4353 presented the canonical residues and is identical to that of profoundly desiccation-tolerant strains,^(28)^ indicating that AB4353 may have this desiccation tolerance phenotype. In fact, as aforementioned, this strain has persisted in HGR clinical settings for, at least, 19 months. Moreover, it has been shown that the copy number of *umuD* and *umuC* error-prone DNA polymerase V genes may directly contribute to desiccation-induced mutagenesis.^(29)^ AB4353 presented one copy of *umuD* and three copies of *umuC*, which may be contributing to increase the mutagenesis rates involved with desiccation-tolerant phenotype.

### Genomic characterization of the pyomelanin biosynthetic pathway

Pyomelanin biosynthetic pathway is well known in *Pseudomonas* species.^(13,14,18,20)^ However, although the pyomelanin production had already been demonstrated in *A. baumannii*,^(15)^ its biosynthesis remains to be characterized in this species. Thus, we performed comparative genomic analysis to identify and characterize the genes involved with pyomelanin production in AB4353. Homolog genes of *hmgR, hmgB* and *hmgC*, involved with pyomelanin central catabolic pathway, were characterized in AB4353, sharing 29%, 45% and 46% deduced amino acid identity with those from *P. putida*, respectively. A homologue of *aroP2* gene, which encodes an aromatic amino acid permease, was found contiguous to the putative *hmgB* in AB4353 (Fig. 2), with a gene arrangement similar to that of *P. putida* KT2440.^(11)^

**Fig. 2:**
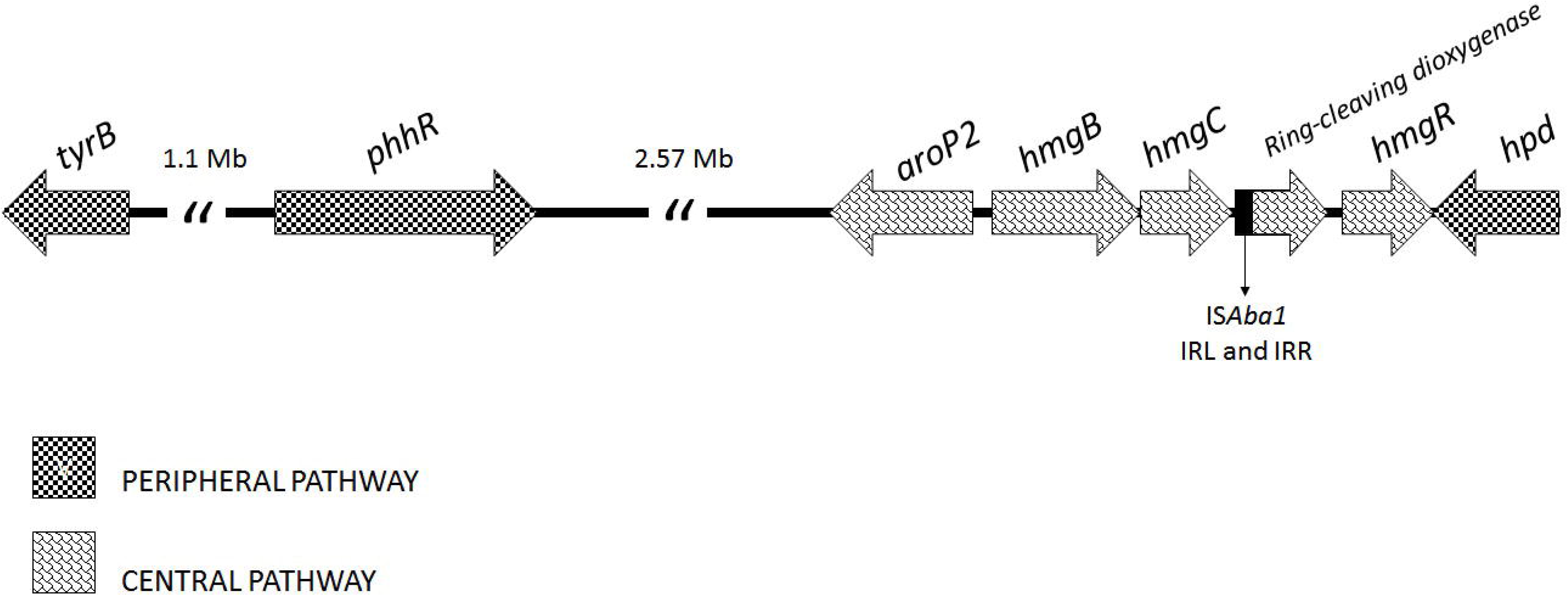
Gene organization of the pyomelanin biosynthetic pathway in *A. baumannii* AB4353. Genes are represented by arrows and the central and peripheral pathways genes are highlighted with different patterns.

The HmgA is a ring-cleaving dioxygenase from the Dioxygenase Superfamily. Although no *hmgA* homologue has been identified in AB4353, a putative gene whose deduced product presented the type I ring-cleaving dioxygenase conserved domain, which is related to the main function of HmgA, was found between *hmgC* and *hmgR* in AB4353 (Fig. 2). Therefore, it suggested the presence of a distant homologue of *hmgA* with no dioxygenase activity in this strain.

The *in silico* analyses revealed that AB4353 harboured the *phhR, hpd* and *tyrB* genes from the peripheral pathway (Fig. 2), which presented 42%, 67% and 45% amino acid identity with those from *P. putida*, respectively. As found for some other genera, the *hpd* and *tyrB* were not linked to the *phh* operon in AB4353, as found for *P. />ulida*In fact, the *phhAB* were absent in AB4353, and the *hpd* was associated with the *hmg* genes (Fig. 2), as observed in *Pseudomonas syringae, Xanthomonas axonopodis, Caulobacter crescentus, Bradyrhizobium japonicum, Mesorhizobium loti*, and *Sinorhizobium meliloti*.^(11)^ Considering that the conversion of phenylalanine in tyrosine is mediated by *phhAB*, and that these genes are absent in AB4353, it can be assumed that a pathway other than hydroxylation of phenylalanine is probably involved in the tyrosine biosynthesis as demonstrated elsewhere.^(11)^

Comparison of the peripheral and central pathways of *Pseudomonas* species and other genera demonstrated a high heterogeneity in gene synteny.^(11)^ In fact, AB4353 displayed a new gene organization concerning both those involved with peripheral and central pathways. Interestingly, a conserved synteny of pyomelanin pathway genes was observed among AB4353 and two other genomes (AB120-02 and AB421) also from IC-5 (ST79) recovered from outbreaks in Spain in 2006-2008 and 2010.^(22,30)^ The unique difference is that in AB421 the *hmgC* was separated from the putative ring-cleaving dioxygenase by a 3.6 kb segment. As aforementioned, the pyomelanin formation depends on the export of the accumulated HGA. AB4353 harboured the entire *hat* ABC transporter gene cluster, sharing 57% (HatA), 69% (HatB), 35% (HatC), 31% (HatD) deduced amino acid identity with those from *P. aeruginosa* UCBPP-PA14.^(14)^

Therefore, i) the presence of the peripheral pathway genes (*phhR, tyrB* and *hpd*) responsible for HGA formation from tyrosine metabolism; ii) the presence of a distant *hmgA* homologue, which is probably not functional, resulting in the HGA cytoplasmic accumulation; and iii) the presence of *hatABCDE* ABC transporter, which pumps HGA, allowing it to self-polymerize into pyomelanin out of the cell; indicate that AB4353 presents the minimum requirements for pyomelanin biosynthesis, and that its production involves a pathway similar to that described in *Pseudomonas* species.^(11,12)^

## CONCLUSIONS

The production of pyomelanin, a pigment associated with virulence in bacteria, by *A. baumannii* strains belonging to the pandemic IC-5, and the existence of a set of genes related to increased adherence and iron uptake, could play a major role in the virulence and persistence of this pandemic lineage.

## ACKNOWLEDGMENTS

We acknowledge Dr. Beatriz Moreira for kindly provided the pyomelanin-producing *A. baumannii* 456MDp used as control in our study.

## AUTHOR CONTRIBUTIONS

EF – Conceptualization and design of the study, performed the experiments, analyzed and interpreted the data, wrote, reviewed and edited the manuscript; RC – collected the bacterial strains; FF and SM – performed the experiments; ACV – Conceptualization and design of the study, scientific supervision, funding acquisition, revision, edition and final approve the manuscript. All authors have read and agreed to the published version of the manuscript.

## REFERENCES

1. Rocha IV, Xavier DE, Almeida KRH, Oliveira SR, Lea NC. Multidrug-resistant *Acinetobacter baumannii* clones persist on hospital inanimate surfaces. Braz J Infect Dis. 2018; 22: 438–41. doi: 10.1016/j.bjid.2018.08.004.

2. Zarrilli R, Pournaras S, Giannouli M, Tsakris, A. Global evolution of multidrug-resistant *Acinetobacter baumannii* clonal lineages. Int J Antimicrob Agents. 2013; 41: 11–19. doi: 10.1016/j.ijantimicag.2012.09.008.

3. Karah N, Sundsfjord A, Towner K, Samuelsen, Ø. Insights into the global molecular epidemiology of carbapenem non-susceptible clones of *Acinetobacter baumannii*. Drug Resist Updat. 2012;15: 237–47. doi: 10.1016/j.drup.2012.06.001.

4. Jones CL, Clancy M, Honnold C, Singh S, Snesrud E, Onmus-Leone F, et al. Fatal outbreak of an emerging clone of extensively drug-resistant *Acinetobacter baumannii* with enhanced virulence. Clin Infect Dis. 2015; 61: 145–54. doi: 10.1093/cid/civ225.

5. McConnell MJ, Actis L, Pachón J. *Acinetobacter baumannii:* human infections, factors contributing to pathogenesis and animal models. FEMS Microbiol Rev. 2013; 37: 130–55. doi: 10.1111/j.1574-6976.2012.00344.x.

6. Giannouli M, Antunes LC, Marchetti V, Triassi M, Visca P, Zarrilli, R. Virulence-related traits of epidemic *Acinetobacter baumannii* strains belonging to the international clonal lineages I-III and to the emerging genotypes ST25 and ST78. BMC Infect Dis. 2013; 13: 282. doi: 10.1186/1471-2334-13-282.

7. Nosanchuk JD, Casadevall A. The contribution of melanin to microbial pathogenesis. Cell Microbiol. 2003;5:203–223. doi: 10.1046/j.1462-5814.2003.00268.x.

8. Yabuuchi E, Ohyama A. Characterization of “Pyomelanin”-Producing Strains of *Pseudomonas aeruginosa*. Int J Syst Bacteriol. 1972;22:53–64.

9. Chatfield CH, Cianciotto NP. The secreted pyomelanin pigment of *Legionella pneumophila* confers ferric reductase activity. Infect Immun. 2007;75:4062–4070. doi: 10.1128/IAI.00489-07.

10. Valeru SP, Rompikuntal PK, Ishikawa T, Vaitkevicius K, Sjöling A, Dolganov N, et al. Role of melanin pigment in expression of *Vibrio cholerae* virulence factors. Infect Immun. 2007; 77: 935–42. doi: 10.1128/IAI.00929-08.

11. Arias-Barrau E, Olivera ER, Luengo JM, Fernández C, Galán B, García JL, et al. The homogentisate pathway: a central catabolic pathway involved in the degradation of L-phenylalanine, L-tyrosine, and 3-hydroxyphenylacetate in *Pseudomonas putida*. J Bacteriol. 2004; 186: 5062–77. doi: 10.1128/JB.186.15.5062-5077.2004.

12. Rodríguez-Rojas A, Mena A, Martín S, Borrell N, Oliver A, Blázquez J. Inactivation of the *hmgA* gene of *Pseudomonas aeruginosa* leads to pyomelanin hyperproduction, stress resistance and increased persistence in chronic lung infection. Microbiology 2009;155:1050–1057. doi: 10.1099/mic.0.024745-0.

13. Ranjan VK, Saha T, Mukherjee S, Chakraborty R. Draft Genome Sequence of a Novel Bacterium, *Pseudomonas sp*. Strain MR 02, Capable of Pyomelanin Production, Isolated from the Mahananda River at Siliguri, West Bengal, India. Genome Announc. 2018; 6: e01443–17. doi: 10.1128/genomeA.01443-17.

14. Hunter RC, Newman DK. A putative ABC transporter, *hatABCDE*, is among molecular determinants of pyomelanin production in *Pseudomonas aeruginosa*. J Bacteriol. 2010;192:5962–5971. doi: 10.1128/JB.01021-10.

15. Coelho-Souza T, Martins N, Maia F, Frases S, Bonelli RR, Riley LW, et al. Pyomelanin production: a rare phenotype in *Acinetobacter baumannii*. J Med Microbiol. 2014;63:152–154. doi: 10.1099/jmm.0.064089-0.

16. CLSI. Performance standards for antimicrobial susceptibility testing: Twenty-second informational supplement M100-S27. Wayne, PA, USA: CLSI; 2017.

17. Magiorakos AP, Srinivasan A, Carey RB, Carmeli Y, Falagas ME, Giske CG, et al. Multidrug-resistant, extensively drug-resistant and pandrug-resistant bacteria: an international expert proposal for interim standard definitions for acquired resistance. Clin Microbiol Infect. 2012; 18:268–281. doi: 10.1111/j.1469-0691.2011.03570.x.

18. Arai T, Hamajima H, Kuwahara S. Pyomelanin production by *Pseudomonas aeruginosa*. I. Transformation of pyomelanin productivity. Microbiol Immunol. 1980;24:1–10.

19. Turick CE, Caccavo F Jr, Tisa LS. Pyomelanin is produced by *Shewanella algae* BrY and affected by exogenous iron. Can J Microbiol. 2008;54:334–339. doi: 10.1139/w08-014.

20. Zeng Z, Cai X, Wang P, Guo Y, Liu X, Li B, et al. Biofilm Formation and Heat Stress Induce Pyomelanin Production in Deep-Sea *Pseudoalteromonas sp*. SM9913. Front Microbiol. 2017;8:1822. doi: 10.3389/fmicb.2017.01822.

21. Caldart RV, Fonseca EL, Freitas F, Rocha L, Vicente AC. *Acinetobacter baumannii* infections in Amazon Region driven by extensively drug resistant international clones, 2016-2018. Mem Inst Oswaldo Cruz. 2019;114:e190232. doi: 10.1590/0074-02760190232.

22. Pérez A, Merino M, Rumbo-Feal S, Álvarez-Fraga L, Vallejo JA, Beceiro A, et al. The FhaB/FhaC two-partner secretion system is involved in adhesion of *Acinetobacter baumannii* AbH12O-A2 strain. Virulence. 2017;8:959–974. doi:10.1080/21505594.2016.1262313.

23. Melvin JA, Scheller EV, Noël CR, Cotter PA. New Insight into Filamentous Hemagglutinin Secretion Reveals a Role for Full-Length FhaB in *Bordetella* Virulence. mBio. 2015;6 pii: e01189–15.

24. Loehfelm TW, Luke NR, Campagnari AA. Identification and characterization of an *Acinetobacter baumannii* biofilm-associated protein. J Bacteriol. 2008;190:1036–1044. doi: 10.1128/JB.01416-07.

25. Tomaras AP, Dorsey CW, Edelmann RE, Actis LA. Attachment to and biofilm formation on abiotic surfaces by *Acinetobacter baumannii:* involvement of a novel chaperone-usher pili assembly system. Microbiology. 2003;149:3473–3484.doi: 10.1099/mic.0.26541-0.

26. Brossard KA, Campagnari AA. The *Acinetobacter baumannii* biofilm-associated protein plays a role in adherence to human epithelial cells. Infect Immun. 2012;80:228–233. doi: 10.1128/IAI.05913-11.

27. Mihara K, Tanabe T, Yamakawa Y, Funahashi T, Nakao H, Narimatsu S, et al. Identification and transcriptional organization of a gene cluster involved in biosynthesis and transport of acinetobactin, a siderophore produced by *Acinetobacter baumannii* ATCC 19606T. Microbiology. 2004;150:2587–2597. doi: 10.1099/mic.0.27141-0.

28. Farrow JM, Wells G, Pesci EC. Desiccation tolerance in *Acinetobacter baumannii* is mediated by the two-component response regulator BfmR. PLoS One. 2018;13:e0205638. doi: 10.1371/journal.pone.0205638.

29. Norton MD, Spilkia AJ, Godoy VG. Antibiotic resistance acquired through a DNA damage-inducible response in *Acinetobacter baumannii*. J Bacteriol. 2013;195:1335–1345. doi: 10.1128/JB.02176-12.

30. Merino M, Alvarez-Fraga L, Gómez MJ, Aransay AM, Lavín JL, Chaves F, et al. Complete Genome Sequence of the Multiresistant *Acinetobacter baumannii* Strain AbH12O-A2, Isolated during a Large Outbreak in Spain. Genome Announc. 2014;2:e01182–14. doi: 10.1128/genomeA.01182-14.

